# The heparan sulfate proteoglycan Syndecan-1 influences local bone cell communication via the RANKL/OPG axis

**DOI:** 10.1101/852590

**Authors:** Melanie Timmen, Heriburg Hidding, Martin Götte, Thaqif El Khassawna, Daniel Kronenberg, Richard Stange

## Abstract

The heparan sulfate proteoglycan Syndecan-1, a mediator of signals between the extracellular matrix and cells involved is able to interact with OPG, one of the major regulators of osteoclastogenesis. The potential of osteoblasts to induce osteoclastogenesis is characterized by a switch of OPG (low osteoclastogenic potential) towards RANKL production (high osteoclastogenic potential).

In the present study, we investigated the influence of endogenous Syndecan-1 on local bone-cell-communication via the RANKL/OPG-axis in murine osteoblasts and osteoclasts in wild type and Syndecan-1 lacking cells. Syndecan-1 expression and secretion was increased in osteoblasts with high osteoclastogenic potential. Syndecan-1 deficiency led to increased OPG release by osteoblasts that decreased the availability of RANKL. In co-cultures of Syndecan-1 deficient osteoblasts with osteoclast these increased OPG in supernatant caused decreased development of osteoclasts. Syndecan-1 and RANKL level were increased in serum of aged WT mice, whereas Syndecan-1 deficient mice showed high serum OPG concentration. However, bone structure of Syndecan-1 deficient mice was not different compared to wild type.

In conclusion, Syndecan-1 could be regarded as a new modulator of bone-cell-communication via RANKL/OPG axis. This might be of high impact during bone regeneration or bone diseases like cancer where Syndecan-1 expression is known to be even more prevalent.

## Introduction

Syndecans, a family of four heparan sulfate proteoglycans (HSPG), are known to be expressed in many different cell types at low levels, but also in a differentiation dependent manner ^1^. Syndecans are composed of an extracellular domain that contains up to three binding sites for GAG chains of different types, e.g. HS chains, some Syndecans also have chondroitin (CS) side chains, near the transmembrane domain ^4^. Additionally, the GAG chain composition is tissue specific, which influences the interaction with a high variety of ligands ^5,6^. The extracellular domain can be shed from the cell surface at a well-characterized site close to the transmembrane region. Intracellularly, Syndecan proteins show a small cytoplasmic domain consisting of a variable domain flanked of two conserved domains that are responsible for intracellular signalling and interaction with the cytoskeleton. These proteins were shown to be up regulated during pathological processes (e.g. inflammation, cancer, healing processes, infection) and seemed to have an important function there. In different models of diseases deficiency of Syndecan-1 led to severe phenotypes with higher mortality or more severe progression of many diseases (see review: Teng *et al*. ^2^), but no obvious phenotype could be observed in an unchallenged organisms lacking Syndecan-1. In previous studies, we characterized Syndecan-4 deficient mice that were not showing any obvious bone phenotype, but we were able to reveal an important function of Syndecan-4 in chondrocytes during enchondral ossification and fracture healing ^3^.

Recently, some studies have drawn additional attention to heparan sulfate proteoglycans (HSPG) as new regulatory players of bone cell communication. Osteoblasts and osteoclasts are known to mediate bone formation and bone resorption via RANKL/OPG balance ^7^. In contrast to RANKL, OPG has a heparin binding domain and was shown to have a high affinity for GAGs ^8^. It has been shown that isolated glycosaminoglycans (GAG) or heparin molecules stimulated osteoclastogenesis, and long term treatment with heparin lead to bone loss ^9,10^. Proteoglycans were found in one complex together with RANKL/RANK and OPG, but heparin molecules also inhibited the binding of OPG to RANKL ^8,9,12^. HS was identified as a component that caused a conformational change in OPG upon binding followed by dimerization of OPG which then resulted in an increased affinity of OPG to RANKL ^17^. Additionally, the immobilization of OPG at the cell surface via binding to membrane bound HSPG seemed to facilitate the interaction with membrane bound RANKL, and this inhibited osteoclastogenesis ^18^. In a recent study using knock-in mice and MC3T3 osteoblastic cell line with an OPG mutant that was incapable of heparan sulfate binding, the interaction of OPG with HSPG on the surface of osteoblasts was indispensable for normal bone homeostasis ^19^.

Especially in bone tumour development and metastasis, the interaction of proteoglycans like Syndecan-1 with OPG was characterised^11^. Sanderson *et al*. and Velasco *et al*. have summarized the results of several studies investigating the role of proteoglycans and Syndecan-1 during osteolytic tumour growth and metastasis ^5,12^. Syndecan-1 was upregulated, either membrane bound or as a soluble form, during a variety of cancers ^6,13,14^. The protein has been recognized as a specific HSPG to influence osteoclastogenesis via OPG binding in bone metastases of multiple myeloma. Behnad-Mehner *et al*. targeted Syndecan-1 expression in MCF-7 breast cancer cells and found ansubsequent increase of OPG leading to inhibition of osteoclastogenesis ^11^.

Because of its interaction with OPG, Syndecan-1 seems to be a potential candidate to influence bone cell communication and bone remodelling also under healthy conditions (bone development, bone homeostasis, aging). The overall question of this study was if Syndecan-1 influences bone cell communication via the OPG/RANKL axis leading to changes in bone structure in healthy organisms. This was investigated by using osteoblasts and osteoclasts isolated from unchallenged WT and Syndecan-1 deficient mice. The capacity of bone cell communication using co-culture *in vitro* was analysed as well as the impact of Syndecan-1 deficiency on bone structure and Syndecan-1 appearance during aging of mice.

## Results

### Syndecan-1 is expressed in osteoblasts and osteoclasts

Expression of Syndecan-1-4 in osteoblasts and osteoclasts as well as the impact of Syndecan-1 deficiency on bone cell differentiation were investigated *in vitro* in wild type and Syndecan-1 lacking cells.

During osteoblast differentiation (DM Medium, table 1) mRNA of Syndecan-1 and -3 were expressed at lower levels compared to Syndecan-2 and -4 (Fig. 1 A). In Fig. 1 B, Syndecan-1 protein was detected using antiSyndecan-1 antibody on WT and Syndecan-1 deficient osteoblasts cultured in DM medium for 7 days. WT osteoblasts showed a strong positive staining, whereas Syndecan-1 deficient osteoblasts displayed no detection of Syndecan-1. During differentiation of osteoclasts in presence of rmRANKL, the mRNA expression of *Sdc1* increased significantly (indicated with #) over time and, at day 6, is higher compared to Syndecan-4 (p<0.01), while the expression of other Syndecans remained mostly unchanged (Fig. 1 C). Syndecan-2 is not expressed in osteoclasts under these conditions. This is of importance, as Syndecan-2 has previously been shown to affect RANKL expression in bone marrow cells ^20^. The immunological staining of osteoclasts after 7 days of differentiation showed a positive Syndecan-1 detection in multinucleated cells (Fig. 1 D). Using co-cultured wild type osteoblasts and osteoclasts, after 7 days of culture in DM+ medium (table 1), positive Syndecan-1 detection was shown in osteoblastic cells (OB), in mononucleated as well as in multinucleated osteoclastic cells (OC) (Fig. 1 E). Functional analyses of osteoblasts lacking Syndecan-1 revealed no differences in mineralization capacity of the extracellular matrix compared to wild type cells (Fig. 2 A). Osteoclast differentiation seemed to be slightly retarded in Sdc1-/- cells showing lower numbers of big multinucleated cells, which was also consistent with less resorption (Fig. 2 B).

**Table 1:** Ingredients of culture medium for osteoblasts, osteoclasts and co-cultures

| Medium | Ingredients |
| --- | --- |
| Basal | Alpha-MEM, 10 % FCS, 2 mM L-glutamine penicillin (100 U/ml)/streptomycin (100 µg/ml) |
| DM | Basal plus 0.2 mM L-ascorbic acid 2-phosphate, 10 mM beta-glycerol phosphate, |
| DM+ | DM plus 1 µM PGE2, 10 nM $1,25(\text{OH})_2\text{D}_3$ |

**Figure 1:**
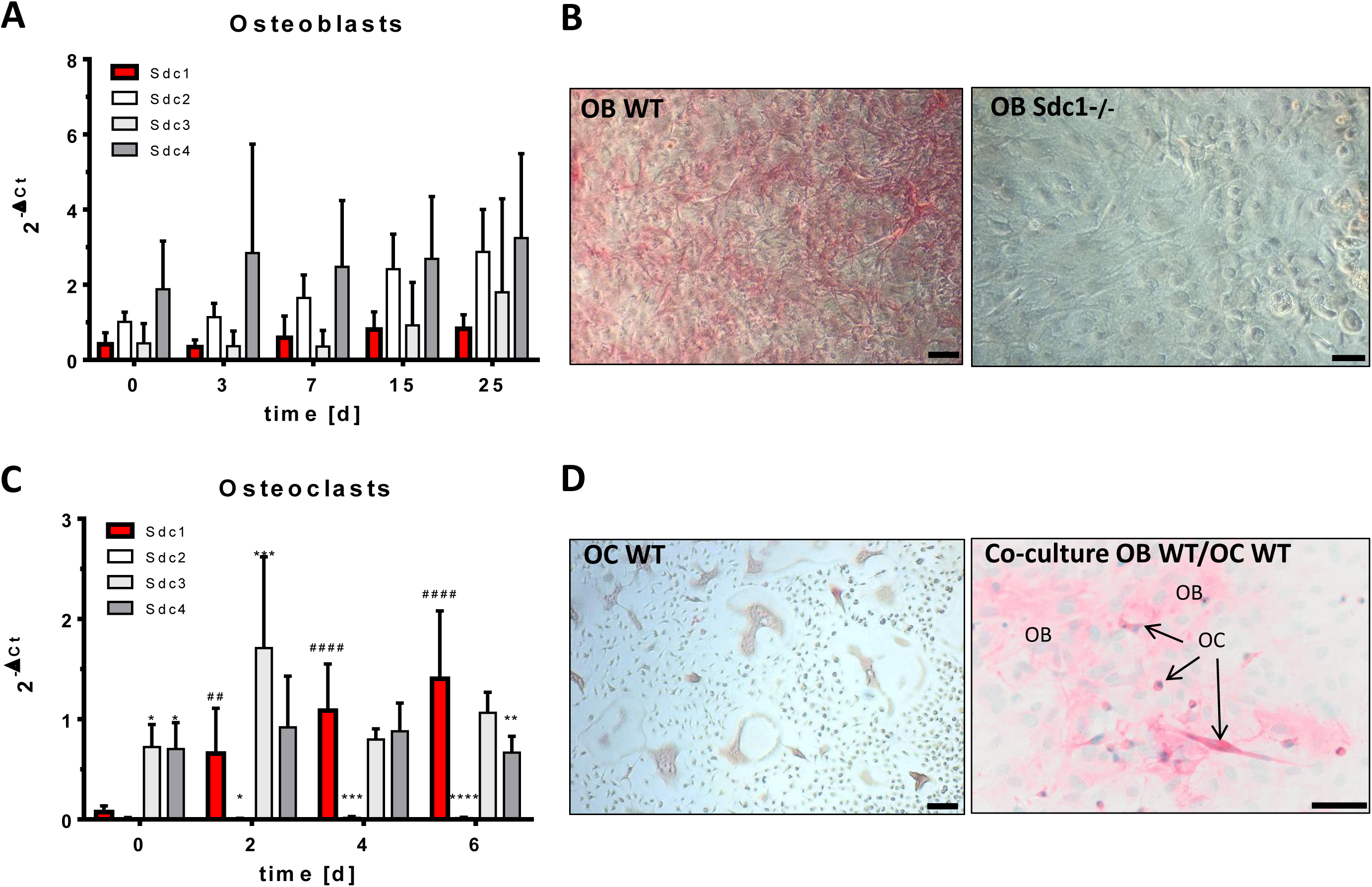
Expression of Syndecans in osteoblasts and osteoclasts during differentiation *in vitro*. Primary osteoblast precursors were isolated from calvariae of newborn mice and seeded in DM-Medium (see Table 1). Osteoclast precursor cells were isolated from bone marrow of long bones of 4-6 week old mice and seeded in DM-Medium supplemented with rmM-CSF and rmRANKL. Co-cultures of osteoblasts and osteoclasts were performed in presence of 1,25(OH)_2_D_3_ and PGE2 (DM+, see Table 1). **A**,**C**: mRNA expression of Syndecan-1-4 (Syndecan-1 in red) were analysed using quantitative real time PCR (Primers see Table 2) normalized to HPRT (ΔCT method). **B, D**: Protein expression of Syndecan-1 in the cells (after 7 days of culture) was detected using a polyclonal antibody against Syndecan-1 (in red) on the cell layer of osteoblasts (OB) in **A**, osteoclasts (OC) in **D** and co-culture of OB (WT) and OC (WT) in **E**. Scale bar: 50µm. Experiments were performed in independent triplicates (A; C). Data are displayed as mean ± SD. AC: 2way ANOVA with Dunnett’s multiple comparison test * indicate expression of Syndecan-1 (red) compared to other Syndecans at each time point, # indicate expression of Syndecan-1 compared to day 0, */#p < 0.05, **/## p<0.01, ***/### p<0.001, ****/#### p<0.0001.

**Figure 2:**
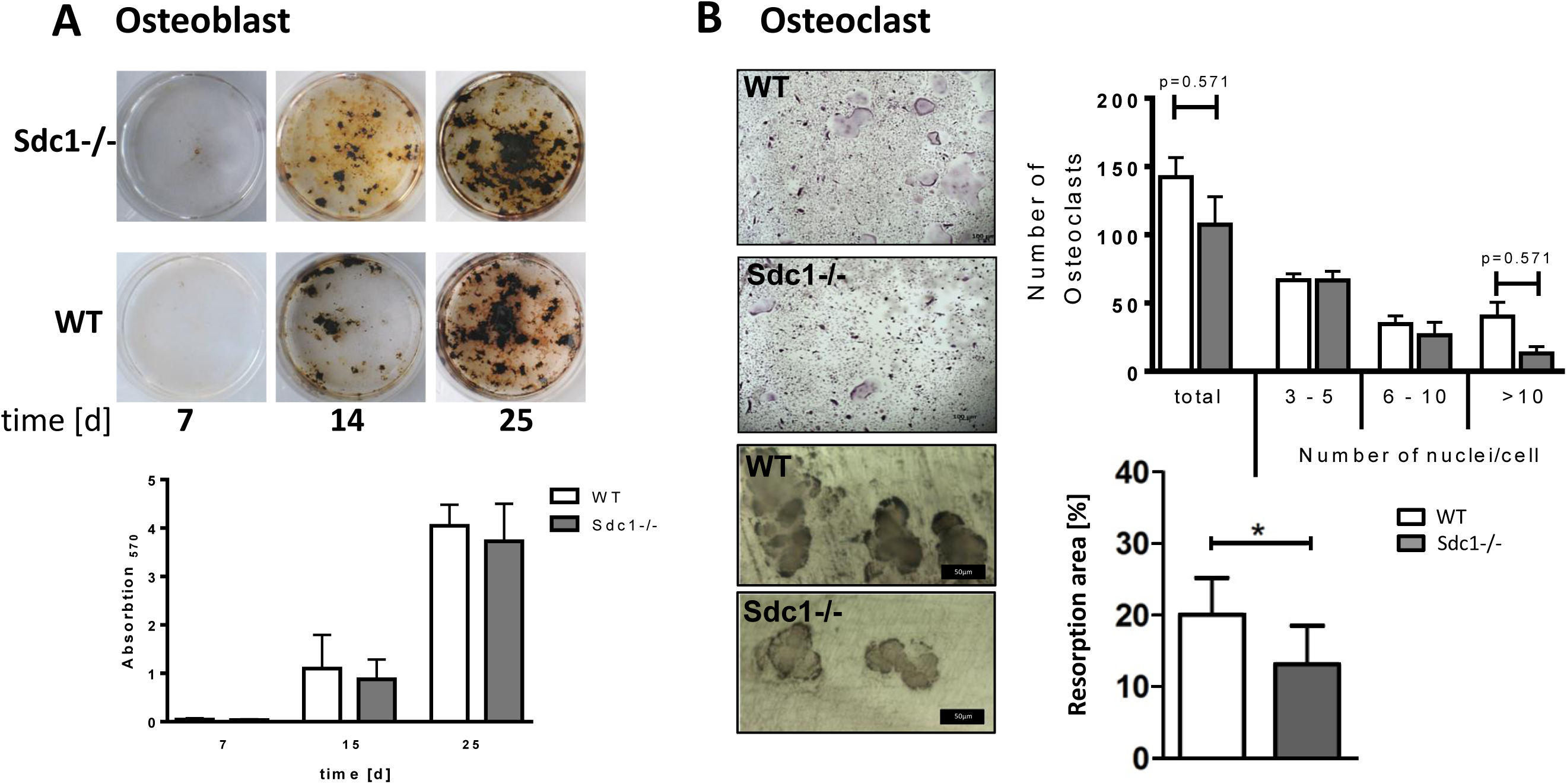
Differentiation and functionality of osteoblasts and osteoclasts dependent on Syndecan-1. **A**: Primary osteoblast precursors (WT, Sdc1-/-) were isolated from calvariae of newborn mice, seeded in DM-Medium (see Table 1) and cultured for up to 25 days. Mineralization of the extracellular matrix was visualized by von Kossa staining and quantified photometrically after alizarin red staining. **B**: Osteoclast precursor cells were isolated from bone marrow of long bones of 4-6 week old mice (WT, Sdc1-/-) and seeded in DM-Medium supplemented with rmM-CSF and rmRANKL. Resorption was analysed using dentin chips. Cells were seeded on cleaned and sterilized dentin chips and cultured in DM-Medium supplemented with rmM-CSF and rmRANKL for up to 9 days. Differentiation of osteoclasts was determined as the development of TRAP positive, multinucleated cells. Resorption was measured as the area of resorption pits per whole area using digital imaging. Experiments were performed in triplicates, scale bars: Trap staining: 100µm, resorption: 50µm, data were presented as mean ± SD, Osteoclast numbers: Kruskal-Wallis test, *p < 0.05, Resorption: Mann-Whitney-U test, * p>0.05.

### Deficiency of Syndecan-1 led to increased OPG release in osteoblasts with high osteoclastogenic potential

We could show, that osteoblastic cells stimulated towards increased osteoclastogenic potential with presence of 1,25(OH)_2_D_3_ and PGE2 (DM+ medium), presented a significant increase in mRNA expression of *Sdc1* (DM: 0.80, DM+: 1.63, p<0.05) (Fig. 3 A) and a slightly higher release of Syndecan-1 protein into the supernatant (Fig. 3 B, DM: 109.5 pg/ml; DM+: 149.5 pg/ml). The amount of Syndecan-1 at the cell surface was not different in presence of DM+ medium compared to DM medium (Fig. 3 C). This was verified by Syndecan-1 immunostaining shown in Fig. 3 D. Because it is known that osteoblasts stimulated with 1,25(OH)_2_D_3_ and PGE2 (DM+) show an induction of RANKL and a reduction of OPG expression, we determined the mRNA expression and release of both proteins into the supernatant of osteoblasts. As shown in Fig. 3 E, RANKL mRNA expression was higher in presence of DM+ medium in both, wild type and Syndecan-1 deficient cells. In contrast to that, OPG mRNA expression was decreased in both genotypes. As expected, release of RANKL protein was increased after induction with DM+ medium, but no RANKL was detectable in the supernatants of Syndecan-1 deficient osteoblasts (Fig. 3 F). It should be noted that our detection system (ELISA) primarily detects free unbound RANKL, thus not complexed with OPG and therefore presumably biological active RANKL. Therefore, in samples with high OPG concentration RANKL is bound to OPG and not detectable, but also not active in RANK interaction. A reduction of OPG concentration in the supernatant of wild type osteoblasts was shown in presence of DM+ medium, but in the supernatant of Syndecan-1 deficient osteoblasts, OPG was only slightly lower compared to DM medium. It was supposed, that immobilization of OPG on the cell surface mediated by HS structures facilitated OPG/RANKL interaction and with that mediates inhibition of osteoclastogenesis. We therefore determined the concentration of RANKL and OPG on the cell surface of WT and Syndecan-1 deficient cells (Fig. 3 G). RANKL could be detected in the cell lysate of WT OB stimulated with DM+ and to a reduced amount in the cell layer of Sdc1-/- cells. In both genotypes, RANKL was increased in presence of DM+ compared to DM. In line with the mRNA expression and the ELISA measurements in the supernatant, OPG in the cell layer was reduced with DM+ medium in WT cells. The same could be seen in Syndecan-1 deficient cells, but comparable to our observations in supernatants, reduction of OPG was less strong. In sum, Syndecan-1 deficient cells show higher amounts of OPG on the cell surface and a higher OPG secretion with DM+ stimulation compared to WT osteoblasts. In contrast to that, less RANKL was bound to the cell surface and no RANKL could be detected in the supernatant of Syndecan-1 deficient cells.

**Table 2:**
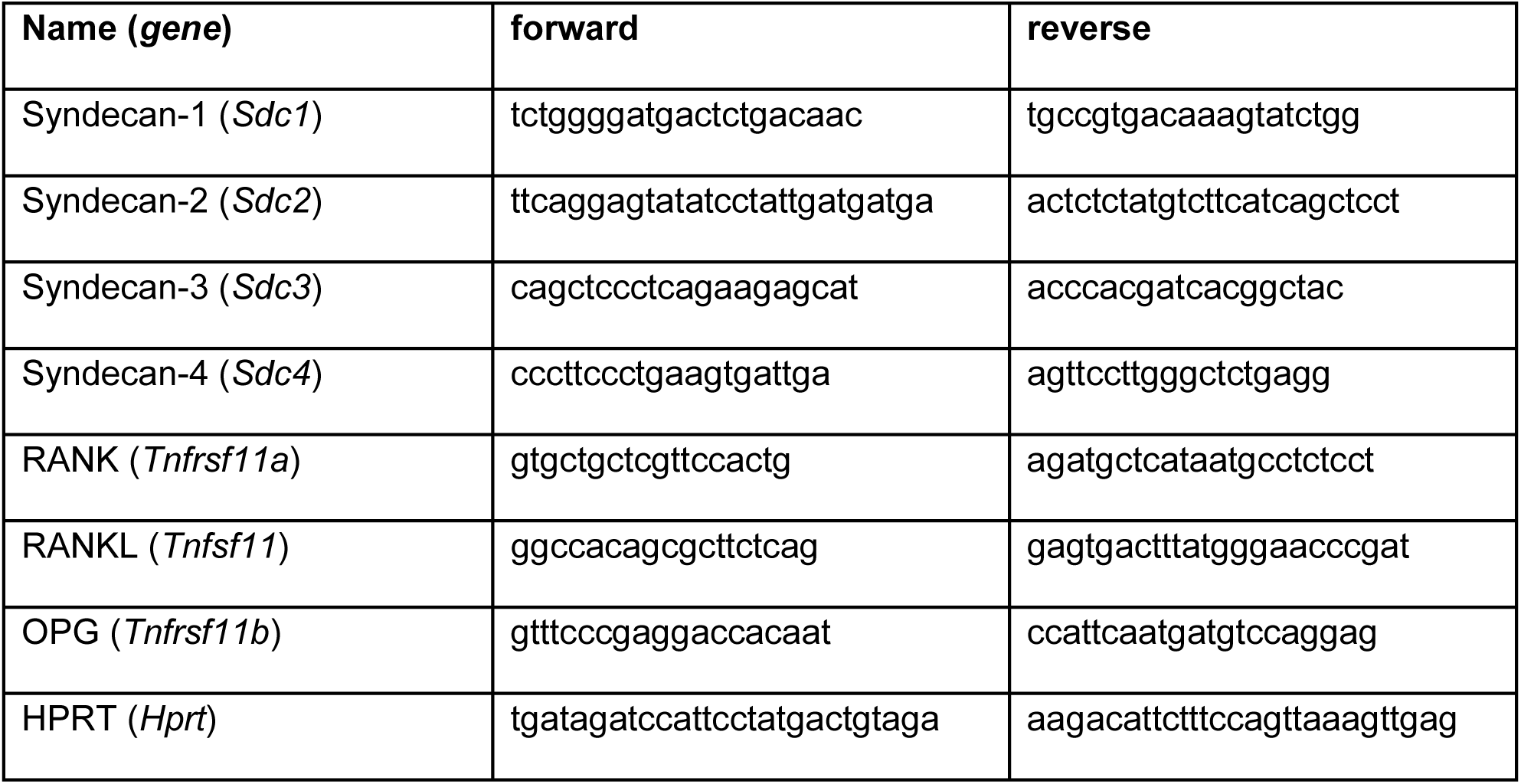
Primers used for quantitative real time PCR

**Figure 3:**
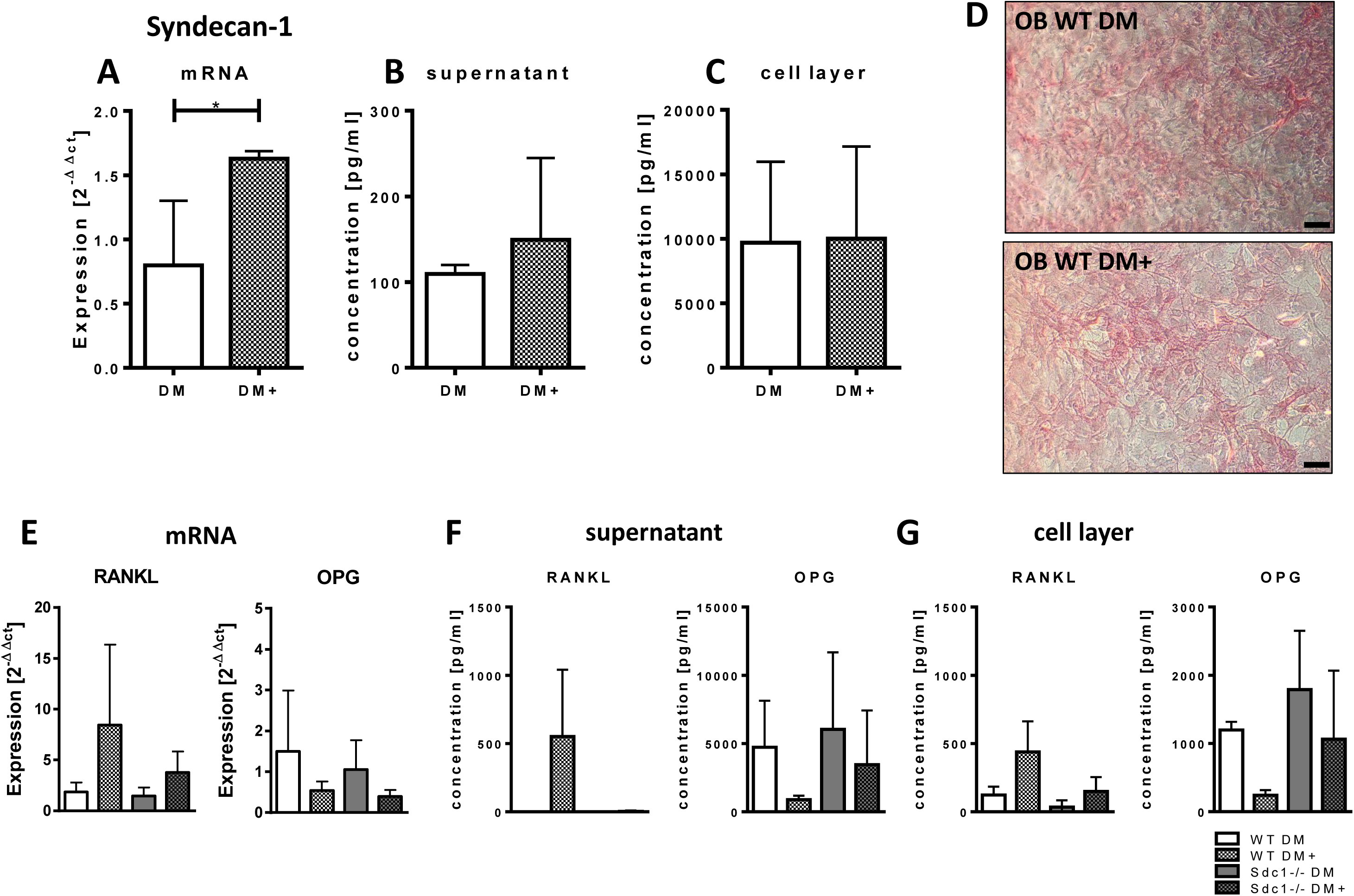
Syndecan-1, RANKL and OPG expression was analysed after stimulation of osteoblastic cells with DM or DM+ medium (day 7). **A**: mRNA expression of Syndecan-1 (in WT cells) was determined by real time PCR. **B, C**: The concentration of soluble Syndecan-1 protein in the supernatant (**B**) and cell layer (**C**) of cells (WT) was analysed by ELISA. Experiments were performed in triplicates. Mann-Whitney-U test, *p < 0.05. **D**: Protein expression of Syndecan-1 in the cells (WT, Sdc1-/-) was detected using a polyclonal antibody against Syndecan-1 (in red) on the cell layer of osteoblasts (OB). Scale bar: 50µm, **E**: mRNA expression of *Tnfsf11* (RANKL) (left) or *Tnfrsf11b* (OPG) (right) in WT and Sdc1-/- osteoblastic cells was determined by real time PCR (Primers see Table 2) normalized to HPRT and day 0 (ΔΔCT method). **F, G**: The concentration of RANKL and OPG protein in the supernatant (**F**) and cell layer (**G**) of cells (WT, Sdc1-/-) was analysed by ELISA. Experiments in E, F were performed in five independent replicates; Experiments in G were performed in duplicates.

#### Increased OPG release in co-cultures of osteoblasts and osteoclasts due to Syndecan-1 deficiency impaired osteoclastogenesis

Bone cell communication via RANKL and OPG under influence of Syndecan-1 was analyzed by differentiation of osteoclast precursors into mature osteoclasts mediated by co-culture of osteoclast precursor cells with RANKL and OPG producing osteoblasts. Stimulation of osteoblasts to increase their osteoclastogenic potential was induced by addition of 1,25(OH)_2_D_3_ and PGE2 (DM+). Co-cultured cells without 1,25(OH)_2_D_3_ and PGE2 showed no osteoclast development. The presence of 1,25(OH)_2_D_3_ and PGE2 alone changed the osteoclastogenic potential of osteoblasts. As shown in figure 4 A, wild type osteoblasts in co-cultures led to the development of mature osteoclasts of both, wild type and Syndecan-1 deficient cells. In co-cultures with Syndecan-1 deficient osteoblasts, significantly decreased osteoclast development was observed (p<0.05, also see quantification in Fig. 4 B). To analyze, if Syndecan-1 deficiency has impact on OPG and RANKL their release was determined in the supernatant of different co-cultured cells (Fig. 4 C). In co-cultures with WT osteoblasts, a high release of RANKL (WT/WT: d3: 216 pg/ml; d7: 620 pg/ml) and low release of OPG (WT/WT: d3: 309 pg/ml, d7: 219 pg/ml) was determined. In comparison to that, co-cultures with Syndecan-1 deficient osteoblasts displayed lower amounts of RANKL (Sdc1-/-/WT: d3: 81 pg/ml, p<0.05; d7: 309 pg/ml, p<0.0001) and high amounts of OPG release (Sdc1-/-/WT: d3: 1930 pg/ml, p<0.001; d7: 1948 pg/ml, p<0.001) into the supernatants. Again, detection of RANKL in complex with OPG by ELISA is prevented, concluding that although RANKL mRNA expression is increased low amounts of active RANKL could be detected. The difference is further illustrated by the calculation of the RANKL/OPG ratio in figure 4 D showing a shift in the balance from RANKL towards OPG in cultures with Syndecan-1 deficient osteoblasts that was highly significant at day 7 of the culture (p<0.001). mRNA expression studies of the corresponding genes of RANK, RANKL and OPG revealed no striking differences between the genotypes or cell types cultivated separately or in co-culture (see supplemental figure 1). In sum, modulation of Syndecan-1 acts on protein level and not on transcriptional regulation.

**Figure 4:**
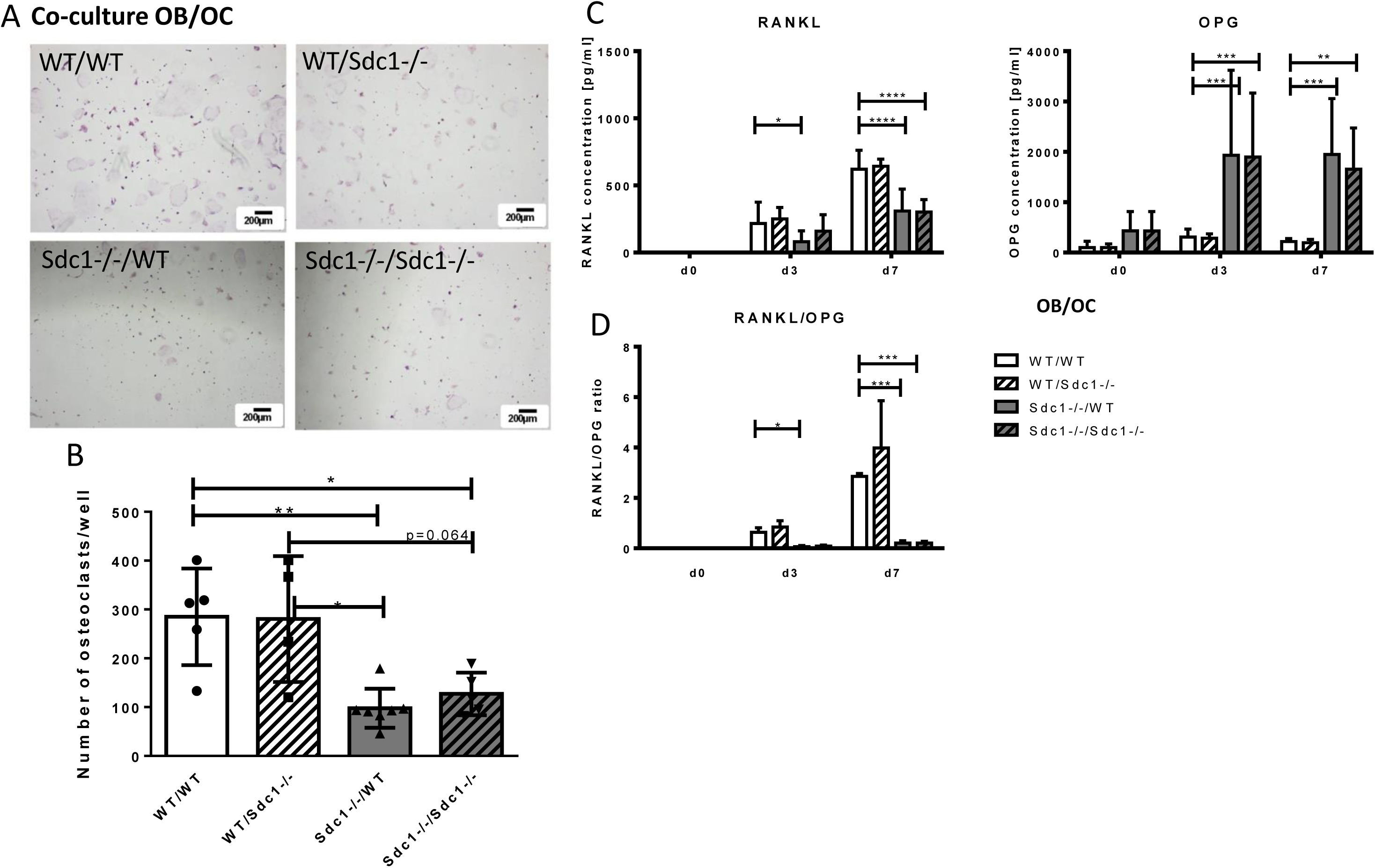
Osteoclast differentiation and RANKL/OPG release in co-cultures of osteoblasts and osteoclasts using Syndecan-1 deficient cells. **A, B**: Primary osteoblast and osteoclast precursor cells were cultured together in DM+-medium for up to 7 days. Different combinations of genotypes (WT/Sdc1-/-) as depicted were used (OB/OC). After 7 days, the osteoblast layer was removed by pulling and TRAP staining was performed, representative images are shown (**A**, scale bar 200µm). Osteoclast number was determined as TRAP positive, multinucleated cells (**B**), Experiments were performed four or more independent replicates, data represent means ± SD, Mann-Whitney-U-test, *p < 0.05, **p<0.01. C: RANKL and OPG concentrations were determined in cell culture supernatant using ELISA. D: RANKL/OPG ratio was calculated. Experiments were performed in independent triplicates, data represent mean ± SD, 2way ANOVA with Bonferroni’s test, *p < 0.05, **p<0.01, ***p<0.001

#### Serum Syndecan-1 and OPG level were increased in mice during aging

Syndecan-1 is already used as a serum marker that is increased during various diseases, but has not been analysed in the course of aging in mice or humans before. We therefore determined the concentration of soluble Syndecan-1 (CD138) in serum of healthy 4, 12 and 18 month old mice. As shown in figure 5 A, in the serum of 7 from 9 young mice no Syndecan-1 was detected. Two serum samples showed a positive detection with a concentration of 524 pg/ml and 734 pg/ml of Syndecan-1. The number of Syndecan-1 positive serum as well as the maximum values of Syndecan-1 concentrations increased with age. In serum of 12 month old mice, 9 of 13 samples were positive for Syndecan-1 with a mean of 384 pg/ml, samples showed a maximal value 2094 pg/ml. Almost all 18 month old mice showed Syndecan-1 positive detection (8 of 9 samples) with highest mean value of 829 pg/ml and maximal values in three samples of up to 2274 pg/ml.

**Figure 5:**
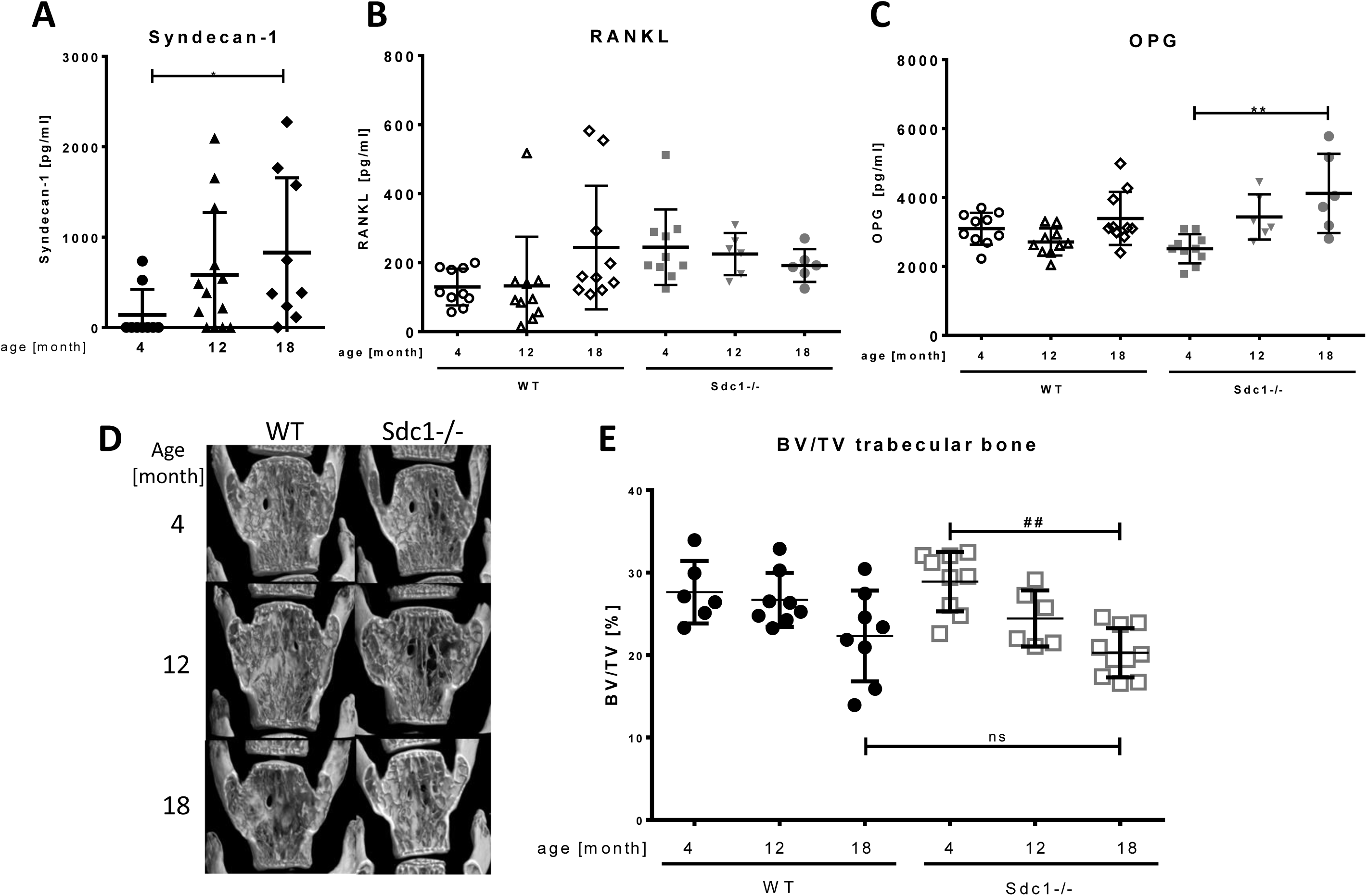
Influence of Syndecan-1 deficiency on serum concentration of RANKL and OPG and bone structure in mice during aging. **A, B, C**: Serum was collected from female mice (WT (all), Syndecan-1 KO (only in **B**,**C**)) of 4, 12 and 18 month of age and concentration of Syndecan-1 (**A**), RANKL (**B**) and OPG (**C**) was determined by ELISA. Data are displayed as mean ± SD, Kruskal-Wallis test with Dunn’s post hoc test, n =9-13 per group. *p < 0.05, **p<0.01. D, **E**: Lumbar vertebrae of mice (wild type, Syndecan-1-/-) at the age of 4, 12 and 18 month were isolated and fixed using 4% PFA. Bone structure was assessed by µCT. Representative pictures of fifth lumbar vertebrae of each age group are shown in **D**, Quantitative analysis of bone volume/tissue volume (BV/TV) of the trabecular bone in the vertebra body is shown in **E**. See Supplemental table 1 for additional bone parameters. Data are displayed as mean ± SD, Kruskal-Wallis test with Dunn’s post hoc test, n =6-10 per group. ^#^p < 0.05, ^##^p<0.01, ns: no significant difference.

Because the balance of osteoblast and osteoclast differentiation is dependent on the ratio of RANKL and OPG, we analysed the amount of these cell communication molecules in the serum of WT and Syndecan-1 -/- mice of different age (n>6). As shown in figure 5 B, the values of soluble RANKL were low (129 ± 53 pg/ml) in young wild type mice and nearly doubled in aged mice (243 ± 179 pg/ml). Overall OPG levels were around 10 times higher than RANKL and did not change significantly in wild type mice during aging (means: 2710-3392 pg/ml, see Fig. 5 C). Young Syndecan-1 deficient mice however showed higher concentration of RANKL (245 ± 110 pg/ml) compared to WT mice (129 ± 53 pg/ml), but these values did not vary during aging (225 ± 61 pg/ml in serum of 12 and 192 ± 48 pg/ml in 18 month old mice). In contrast to WT mice, OPG levels were low in young Syndecan-1-/- mice (2513 ± 423 pg/ml), but increased significantly with aging (12 month: 3437 ± 656 pg/ml; 18 month: 4119 ± 1150 pg/ml).

#### Skeletal remodelling of Syndecan-1 deficient mice during bone development and aging was not affected

Analysis of the whole skeleton of newborn mice revealed no difference in bone size or development, appearance of growth plates, bone mineralization or collagen content (Supplemental Fig. 2) between WT and Sdc1-/- mice. Furthermore, using µCT, we analyzed structural parameters of vertebra bones during growth and aging in female Syndecan-1 deficient mice and age matched wild type animals. As shown in figure 5 D, E, the ratio trabecular bone volume/tissue volume (BV/TV) of lumbar vertebra of Syndecan-1 deficient mice of all age groups showed decreasing values for bone mass like in wild type mice during aging. Overall, no significant differences were found between genotypes. Further parameters of bone structure (whole BV/TV, cortical thickness, trabecular number, trabecular separation, trabecular thickness) were determined as well, but also showed no significant differences between age matched genotypes (Supplemental table 1).

## Discussion

Syndecan-1 was shown to bind OPG via its heparan sulfate proteoglycan side chains and thereby influences bone resorption during breast cancer and multiple myeloma ^11,15^. However, little is known about their function under physiological conditions when the organism is unchallenged. Therefore, we investigated the impact of Syndecan-1 and its deficiency on osteoblast-osteoclast communication.

We first characterized expression and regulation of Syndecan-1 in osteoblasts and osteoclasts and its role in direct bone cell communication. Expression analysis of osteoclasts revealed specific upregulation of Syndecan-1 during differentiation, and a slight retardation of differentiation and bone resorbing capacity in Syndecan-1 deficient osteoclastic cells. Sdc1-/- osteoblasts were shown to mineralize their extracellular matrix in a comparable way like wild type cells. Without any stimulation, Syndecan-1 as well as Syndecan-2, -3 and -4 were expressed in these cells at comparable and low levels. From these results, we have concluded, that osteoblasts as well as osteoclasts themselves were not functionally impaired by the lack of Syndecan-1 although osteoclast development was slightly retarded. Next, we investigated the communication capacity of osteoblasts and osteoclasts with regard to osteoblast driven osteoclast stimulation by OPG and RANKL referred to as osteoclastogenic potential of the osteoblast. Firstly, we could show, that Syndecan-1 expression and protein production in osteoblasts was dependent on culture conditions using 1,25(OH)_2_D_3_ /PGE2 (DM+) as a stimulus of osteoclastogenic potential in osteoblasts. Besides an increase of RANKL and a repression of OPG, which was expected, Syndecan-1 expression and release into the supernatant was induced in presence of DM+. Secondly, detection of OPG and RANKL in the supernatants of osteoblasts revealed, that the bioavailability of RANKL in the environment is dependent on OPG and Syndecan-1 protein level. Sdc1-/- osteoblasts stimulated with DM+ medium showed an increased amount of OPG on the cell and as a release into the supernatant, whereas no RANKL could be detected in supernatants of these cells. The presence of Syndecan-1 led to a decreased OPG level on the cell and in the supernatant and free RANKL could be detected in the supernatant. Thirdly, in direct co-cultures of osteoblasts and osteoclasts of different genotypes (WT/Sdc1-/-) under osteoclast inducing conditions (DM+) the presence of Syndecan-1 in osteoblasts was a prerequisite for osteoclast differentiation. This was independent of the genotype of the osteoclasts co-cultured with the osteoblasts. From that, we also concluded that osteoclast derived Syndecan-1 could not compensate for the lack of Syndecan-1 during onset of bone cell interaction most likely due to low expression in osteoclast precursor cells. In line with our experiments with only osteoblasts, a lack of Syndecan-1 in the co-cultured osteoblasts led to a high amount of OPG in the supernatants that was not due to altered OPG mRNA expression. This surplus of OPG did not allow the induction of osteoclastogenesis in these cultures. Because mRNA expression of OPG/RANKL/RANK was not effected by the lack of Syndecan-1, the effect of Syndecan-1 was a post-translational one. We draw the conclusion, that Syndecan-1 on protein level produced by osteoblasts with high osteoclastogenic potential that effected the inhibitory function of OPG on osteoclastogenesis. There are some studies already published that emphasised the role of Syndecan-1 within the interaction of OPG on osteoclastogenesis. Benad-Mehner *et al*. performed knock down experiments in breast cancer cell (MCF7) using siRNA to target Syndecan-1 ^11^. In their experiments, the mRNA expression of OPG was upregulated and more OPG was released into the environment. In indirect co-culture, osteoclastogenesis was impaired as well as resorption. They could rescue the inhibitory effect by OPG neutralizing antibodies. A possible mechanism of OPG/Syndecan-1 interaction was supposed by Standal *et al*. during multiple myeloma ^15^. They postulated that Syndecan-1 expressing myeloma cells bind, internalize and degrade OPG *in vitro* and *in vivo* as concluded from lowered OPG level in serum of patients with multiple myeloma. A study of Kelly *et al*., pointed into a similar direction. They could show that breast cancer cells (MCF-7) that are expressing Syndecan-1 and it’s sheddase Heparanase, induced osteoclastogenesis. They concluded that soluble Syndecan-1 was important for this interaction and, furthermore, they stressed that Syndecan-1 acts as a soluble factor together with other cytokines like IL-8 ^23^. Taken together, this argued to a function of OPG clearance mediated by Syndecan-1, but the localization of Syndecan-1 as a membrane bound or soluble protein remained unclear.

In our setting, the genotype of the osteoblasts determined the fate of osteoclast development. We compared the localization of Syndecan-1, OPG and RANKL on the cells or in the supernatant of osteoblasts with low or high osteoclastogenic potential and found a stronger correlation of function with soluble Syndecan-1 on the RANKL/OPG ratio, but an additional effect of the membrane bound form cannot be excluded. There are strong arguments for a functional important immobilization of OPG on the cell surface of osteoblasts published by Nozawa *et al*., who have shown that OPG bound to HS on the surface of osteoblasts effectively contributed to the inhibition of RANKL interaction to RANK ^18^. Li *et al*. demonstrated that HS on osteoblastic cells determined the conformation of OPG, its dimerization and with that the binding affinity to RANKL ^17^. In both studies, RANKL was considered only as a membrane bound receptor and OPG had to be arranged in a HSPG/OPG/RANKL complex at the osteoblast surface to effectively inhibit RANKL function. However, RANKL is known to be secreted into the environment by many cell types, and even in our co-cultured cells, RANKL can be found as a soluble protein in addition to the surface of osteoblasts. Further investigation is needed to elucidate the impact of Syndecan-1 located on the surface of osteoblasts as well as the role of Syndecan-1 shedding from the surface of the osteoblasts.

From our results and supported by different studies from other groups as discussed above, we concluded that Syndecan-1 on the surface and in the surrounding environment of osteoblastic cells has impact on the interaction of OPG with RANKL during induction of osteoclast differentiation. We hypothesize that the binding of OPG to the HS chains presented by Syndecan-1 interferes with the inhibitory effect of OPG on RANKL and with that supports osteoclastogenesis. In Fig. 6 we propose a model of a previously unknown Syndecan-1 function dependent on the direct interaction of osteoblasts and osteoclasts under anabolic (bone formation) and osteoclastogenic conditions. Bone cell communication during bone formation leads to high OPG expression and release into the environment (Fig. 6 A). Osteoblasts produce some low amount of Syndecan-1, located at the cell surface and released into the environment, and no/little amount of RANKL is released into the environment. Under these conditions, osteoclastogenesis is supressed by an excess of OPG that inactivates RANKL. A shift to osteoclastogenic conditions (Fig. 6 B) is induced by presence of 1,25(OH)_2_D_3_/PGE2, by increase of inflammatory cytokines, stress conditions, aging and many more ^24-27^. As shown in our study, expression of Syndecan-1 is increased in osteoblasts with osteoclastogenic potential. Syndecan-1 molecules block OPG/RANKL interaction and with that promote RANKL stimulation of osteoclastogenesis. This function is lacking if Syndecan-1 is missing (lower panel). Even if OPG expression is downregulated, OPG cannot be removed from the environment and inhibits RANKL and with that osteoclastogenesis.

**Figure 6:**
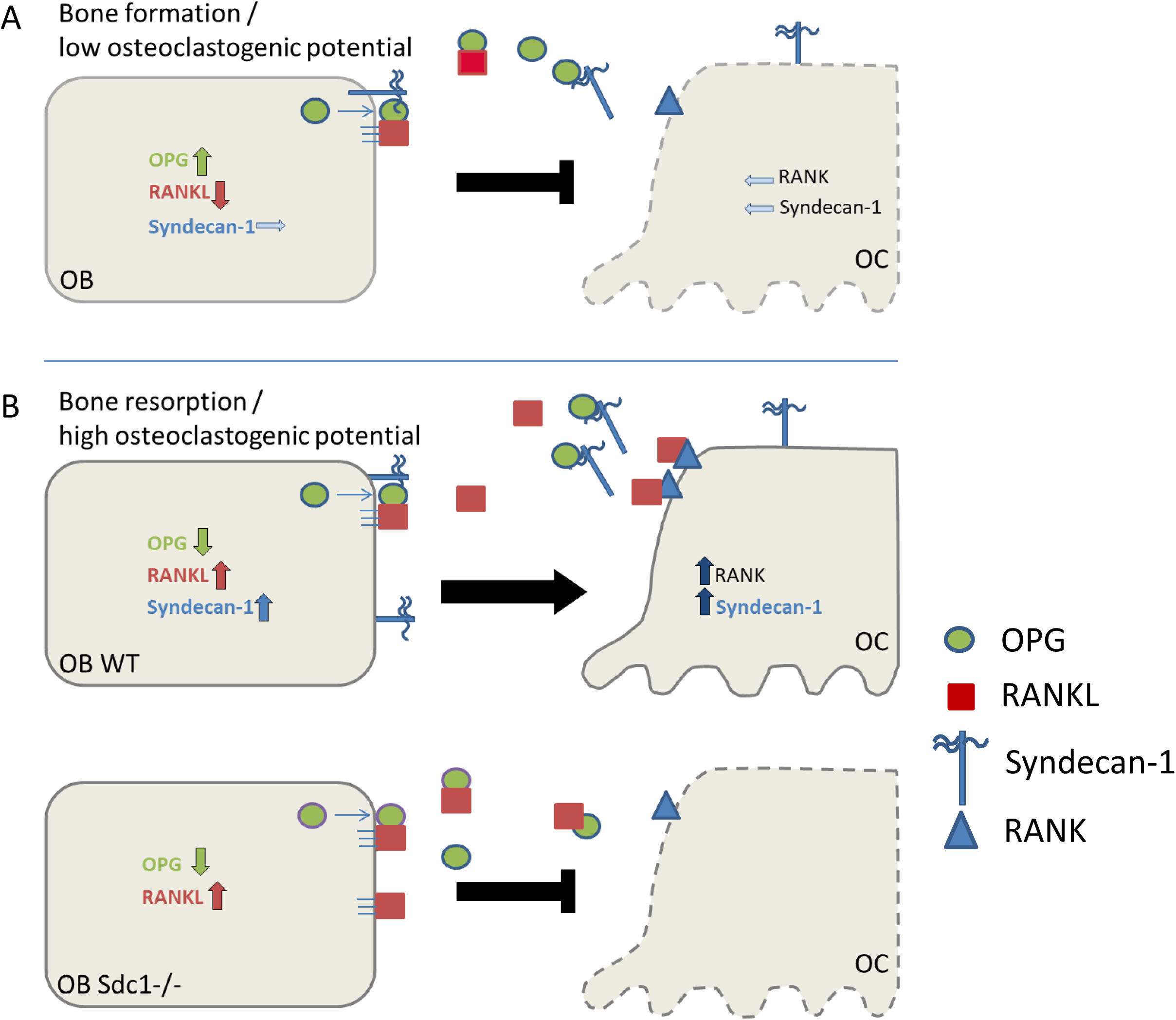
Proposed function of Syndecan-1 in local bone cell communication (model), **A**: Syndecan-1 has a minor function in bone forming osteoblasts with low osteoclastogenic potential. **B**: Syndecan-1 produced by osteoblasts with high osteoclastogenic potential supports osteoclast differentiation by OPG clearance.

To prove a relevance of our findings on Syndecan-1 function in bone cell communication *in vivo*, we determined the amount of RANKL and OPG as well as Syndecan-1 in the serum of mice during aging. Our results revealed an increased release of soluble Syndecan-1 with increasing age of mice. Increasing concentrations of serum Syndecan-1 were also reported in different studies of diseases in humans and mice. In line with our *in vitro* measurements (OB with DM+), RANKL concentrations were increased in aged WT mice whereas OPG remained on the equal levels. Syndecan-1 deficient mice showed increasing OPG levels with age, but RANKL was not elevated in age-matched mice. Investigation of bone development and bone structure during growth and aging by comparing unchallenged wild type and Syndecan-1 deficient mice revealed no significant differences. This is in line with former results that revealed the lack of Syndecan-1 in mice has only minor effects without further stimulus ^22,28,29^. Skeletal morphology and homeostasis is maintained by many different factors like cytokines that influence the RANKL/OPG axis and are produced by different cells besides osteoblasts e.g. activated T-cells or macrophages ^30-32^ but also mechanical stimuli affect bone structure *in vivo*^*33*^ that might mask the effect of Syndecan-1 deficiency in healthy mice during aging. Additionally, as we showed in Fig. 1 A, Syndecan-1 is not the only Syndecan molecule that is produced in osteoblasts, and *in vivo*, additional cell types might contribute to bone cell communication. Possibly, one or more Syndecans compensate for the loss of Syndecan-1 *in vivo*. This is of importance, because for the binding of OPG the presence and sulfatation of HS chains was more important that the core protein. Obvious bone loss was shown only in mice that were deficient for OPG (total or OB specific), lacking the HS binding site in OPG or lacking the *ext1* gene leading to a loss of HS chains ^18,19,34^. Taking into account, that the expression of Syndecan proteins is cell type and differentiation dependent, *in vivo* the pattern of Syndecans on cell surface might rely on environmental stimuli. Regarding Syndecan-1 as a new modulator of bone-cell-communication might be of high impact during bone regeneration or bone diseases where Syndecan-1 expression is known to be even more prevalent.

## Conclusions

Our results provide evidence that Syndecan-1 acts as a new modulator of RANKL/OPG balance by inhibiting OPG binding to RANKL during bone cell communication. This mechanism is based most likely on HS chains carried by Syndecan-1 and OPG within the local microenvironment of bone cells. mRNA expression of OPG or RANKL is independent of Syndecan-1, but osteoclastogenic potential of osteoblast has an impact on Syndecan-1 mRNA expression and protein production in osteoblasts. Further research should reveal if Syndecan-1 has to be localized to the cell surface on either osteoblasts or osteoclasts or if shedding plays a distinct role during this communication process.

## Methods

### Mice

All experiments were performed according to the protocol approved by the Landesamt für Naturschutz, Umweltschutz und Verbraucherschutz (LANUV, AZ: 84-02.05.50.A15.005), Northrhine-Westphalia, Germany according to the legislation for the protection of animals used for scientific purposes (Directive 2010/63/EU). C57Bl/6 Syndecan-1-/- (Sdc1-/-) mice were generated as a global knockout by Stepp *et al*. ^28,35^. Briefly, the signal sequence, exon 1 and a large part of the first intron of the *Sdc1* gene was interrupted by a neomycin resistance cassette driven by a PGK promotor. All mice were inbred homozygously at a C57BL/6 genetic background. Mice were kept under conventional not pathogen-free conditions with a 12 hour light/dark cycle, up to 65% relative humidity and 20°C temperature in open cages grouped with up to 4 animals enriched with nesting material and paper houses. Mice had access to tap water and standard rodent chow (Altromin GmbH, Lage, Germany) ad libitum. Syndecan-1 deficient mice showed no severe phenotype, are vital and fertile.

### Cell culture

For *in vitro* experiments with osteoblasts and osteoclasts, primary precursors of wild type and Syndecan-1 deficient mice were isolated. Primary **osteoblast precursors** were isolated from the calvarias of newborn mice by sequential digestion (0.2 % dispase Grade II (Roche, Sigma-Aldrich, /collagenase Typ Ia, Sigma Aldrich, C-9891). Cells were seeded (2×10^4^ cells/well) in 24-well plates in alphaMEM medium (Merck, Darmstadt, Germany, #F0925) plus 10 % FCS (Gibco, Thermo Fisher, Waltham, MA, USA, #10270), 2 mM L-glutamine (Sigma-Aldrich, Merck #59202C) and penicillin (100 U/ml)/streptomycine (100 µg/ml), supplemented with 0.2 mM L-ascorbic acid 2-phosphate (Sigma-Aldrich, #A8960) and 10 mM beta-glycerol phosphate, Sigma-Aldrich, #G9422; =DM (Differentiation medium for osteoblasts without osteoclastogenic potential, see Table 1) as described previously ^36^. After 24 hours, the non-adherent cells were removed. Isolated osteoblasts were differentiated separately up to 25 days with medium changed every second day. Mineralization was visualized by von Kossa staining and quantified photometrically by Alizarin Red S staining. The **osteoclast precursors** were isolated from the bone marrow of 4-6 week old mice (femur/tibia). For separate culture and differentiation of osteoclasts, precursor cells (1×10^5^ cells/well) were seeded in 96-well plates in DM medium supplemented with rmM-CSF (day 1-3: 50 ng/ml, day 4-7 30 ng/ml, R&D) and rmRANKL (50 ng/ml, R&D). For **co-culture**, cells were seeded (2×10^6^ cells/well) in 6-well plates (alphaMEM plus 10 % FCS, 2 mM L-glutamine and penicillin (100 U/ml)/streptomycin (100 µg/ml) supplemented with 50 ng/ml recombinant murine (rm) M-CSF (R&D, #416-ML). After 24 hours the non-adherent cells were collected, counted and added (4×10^5^ cells/well, 24 well plate) to the adherent osteoblast precursors in presence of DM-medium supplemented with 1 µM PGE2 (Sigma-Aldrich, #P6532) and 10 nM 1,25(OH)_2_D_3_ (Sigma-Aldrich, #D1530) (=DM+; Differentiation medium for osteoblasts with osteoclastogenic potential, see Table 1) ^37^. Under stimulation with 1,25(OH)_2_D_3_ and PGE2 (DM+) osteoblastic cells increase their osteoclastogenic potential by upregulation of RANKL expression and downregulation of OPG expression ^38^. Osteoclasts are located beneath the osteoblasts, that develop a cohesive cell layer. Differentiation of osteoclasts was quantified, in co-cultures after pulling away the osteoblast layer, by counting the TRAP positive, multinucleated cells and nuclei/cell (Acid Phosphatase Leukocyte Kit (387 A), Sigma-Aldrich). Resorption was analysed using dentin chips. Cells were seeded on cleaned and sterilized dentin chips in 96-well (1×10^5^ cells/well) and cultured in DM-Medium supplemented with rmM-CSF and rmRANKL for up to 9 days. Cells were removed after lysis by sonication and resorption pits were stained with Indian ink and visualized using a BX51 microscope (Olympus).

### Detection of RANKL/OPG/Syndecan-1 in supernatant and cell layer and mRNA expression of RANK/RANKL/OPG and Syndecan-1

Concentration of molecules (OPG,RANKL, Syndecan-1) in the supernatant (200µl) and cell layer (1 well, after washing with PBS buffer (Sigma-Aldrich, #D8537-500ML) and lysis using RIPA buffer (150 mM NaCl, 1 % Triton-X-100, 0.5 % Sodiumdeoxycholat, 0.1 % SDS, 50 mM Tris-Base, pH 8,0, protease inhibitor) protein concentration were determined by ELISA (OPG,RANKL: Quantikine ELISA, R&D Systems, Bio-Techne GmbH, Wiesbaden-Norderstedt, Germany, Syndecan-1: Boster Biological Technology). ELISA was performed as instructed by the manufacturer. It should be noted that this ELISA detection system primarily detects free unbound RANKL, thus not complexed with OPG.

### mRNA expression of RANKL/OPG and Syndecan-1

mRNA expression of Syndecan-1, RANKL and OPG was analysed by quantitative real time PCR. RNA was isolated using RNeasy Micro Kit (Qiagen, Hilden, Germany) and the concentration and quality of the RNA were measured photometrically. Reverse transcription of the RNA into cDNA was performed according to the manufactures’ instructions (High-Capacity RNA-to-cDNA Kit, ThermoFischer Scientific). For quantitative real time PCR, primers were used as indicated in Table 2. Besides Syndecan-1, we also determined the expression of the three other Syndecan genes during differentiation of osteoclasts and osteoblasts. HPRT expression was used as an internal housekeeping gene. Relative quantification was performed according to the ΔCT (normalized to HPRT) or ΔΔCT (normalized to HPRT and day 0) method.

### Immunochemistry

Protein synthesis of Syndecan-1 was shown by immunocytochemistry using a polyclonal antibody against murine Syndecan-1 (CD138) clone 281-2 (1:50, rat anti-mouse, BD Pharmingen) in co-cultured cells after fixation with ice-cold 99% methanol and heparitinase III (2.4mIU/ml, Sigma-Aldrich, H2519-50UN)/ chondroitinase ABC Lyase (0.1U/ml, Sigma-Aldrich, C2905) treatment. Amplification and visualization was performed using secondary antibody (1:50, goat anti-Rat IgG, BD Pharmingen, Heidelberg, Germany) conjugated with biotin (visualised by using Vectastein ALP Kit, AK-5000, Vector Laboratories, Burlinggame, CA, USA) and analysed using a BX51 microscope and cell sense dimensions software (Olympus).

### Serum concentrations of Syndecan-1

Serum samples of 4, 12 and 18 month old wild type and Syndecan-1 deficient mice were prepared from blood collected after cervical dislocation. The concentration of Syndecan-1, RANKL and OPG was determined using an ELISA Kit (Syndecan-1: Boster; RANKL/OPG: R&D). In serum of Syndecan-1 deficient mice, no Syndecan-1 was detectable, as expected; therefore, only results of wild type mice are shown.

### Histomorphometry

Histological analysis of the bone phenotype was performed according the nomenclature of Dempster *et al*. using bones of 4, 12 and 18 month old female mice (wild type, Sdc1-/-) as indicated ^39^. Fifth lumbar vertebrae of adult mice (4-18 month) were fixed, kept humid and analysed by µCT (Skyscan 1176, Bruker, Billerica, MA, USA) for mineralized bone structure. µCT scanning and analysis was performed with a voxel resolution of 9 µm according to standard guidelines ^40^. Cortical and trabecular bone were analysed via segmentation using manufactures software (DataViewer, CTvox, CTan). Thresholds for trabecular bone were defined by the Outsu method included in the software.

### Statistics

Statistical analysis was carried out using GraphPad Prism version 6.07 for Windows, GraphPad Software, San Diego, California USA, www.graphpad.com). Different statistical tests (Kruskal-Wallis, Mann-Whitney-U, Holm-Sidak) were used dependent on the groups that are compared and are indicated in the legend of the figures. Results were considered significant with a p-value <0.05.

## Acknowledgements

We thank Simone Niehues, Nina Ehrens and Iska Loessmann for excellent technical assistance. Martin Götte kindly provided the Syndecan-1 deficient mouse strain.

## Authors’ contribution statement

Study design: MT, RS; Study conduct: MT; Data collection: MT, HH, TK; Data analysis: MT, HH, TK; Data interpretation: MT; DK, RS; Drafting manuscript: MT, DK, RS; Revising manuscript content: MG, DK, RS; Approving final version of manuscript: MT, DK, RS; MT takes responsibility for the integrity of the data analysis.

