## Supplemental figures for "The heparan sulfate proteoglycan Syndecan-1 influences local bone cell communication via the RANKL/OPG axis"

**Supplemental Data Legends**

**Supplemental Figure 1**: mRNA expression of RANK, RANKL and OPG in osteoblasts and osteoclasts dependent on Syndecan-1. Primary bone cell precursor cells from mice (WT, Syndecan-1-/-) (OB, OC) were cultured separately or in co-culture (OB/OC). mRNA was isolated after up to 25 days as depicted in the graphs and expression of RANK, RANKL and OPG was determined normalized to HPRT using quantitative real time PCR. Experiments were performed three times, in triplicates, data were presented as mean ± SD. Primer sequences: See table 1 in the manuscript.

- The induction level of RANK over time was much higher (9 to 12 fold) in the co-cultured cells than in separated osteoclastic cells (3 to 4 fold) differentiated in presence of rmRANKL. Possibly the activation of osteoclastognenesis is much more efficient with direct cell cell interaction compared to separated osteoclast culture. RANK expression at low level was observed also in separated primary osteoblast cultures, which point to some contamination with macrophages during preparation that was not further investigated. RANKL expression was low in unstimulated osteoblasts (DM) and absent in osteoclasts single culture. In co-cultures, RANKL expression was increased (4 fold) as shown previously with osteoblasts stimulated with DM+ (Fig. 3 E). OPG gene expression was high, in wild type and Syndecan-1 deficient osteoblasts cultivated in the absence of 1,25(OH)2D3 or PGE2 (lower, left), while, as expected, OPG expression was suppressed under conditions that increase osteoclastogenic potential (lower, right: co-culture, see also Fig. 3 E: single OB).

**Supplemental figure 2**: Bone development in Syndecan-1 deficient mice. A: Whole skeleton of newborn mice were used for alcian blue/alizarin red staining of bone (red) and cartilage (blue), n=6, scale bar 2mm. B: Longitudinal sections of humerus, femur and tibia embedded in paraffin were used for histological quantification of limb development in detail. Maturation of bone development was quantified as ratio of calcified area per whole bone area [%] in the humerus, femur and tibia of wild type and Syndecan-1 deficient mice. C: Paraffin sections of tibia of newborn mice were stained for cartilage (Alcian blue), calcified bone (Masson-Goldner) as well as collagen II and collagen X using primary antibodies and immunohistochemical staining. Data were presented as mean ± SD.

- No differences were found.

**Supplemental table 1**: Bone parameters of wild type and Syndecan-1 deficient mice during aging
Bone structure was assessed by µCT in vertebra of 4 to 18 month old female Syndecan-1 deficient and wild type mice. Data are presented as mean ± SD, n = 6–10 per group. TV = Tissue volume; BV/TV = bone volume fraction; Tb.N = trabecular number; Tb.Th = trabecular thickness; Tb.Sp = trabecular separation; Ct.Th = cortical thickness. Kruskal-Wallis test with Dunn´s post hoc test, n =9-13 per group. * indicate significant difference within age matched genotypes: no significant differences found, # indicate significant difference to 4 month old group of the same genotype, #p>0.05, ##p>0.01, ###p>0.001, ####p>0.0001

Supplemental figure 1

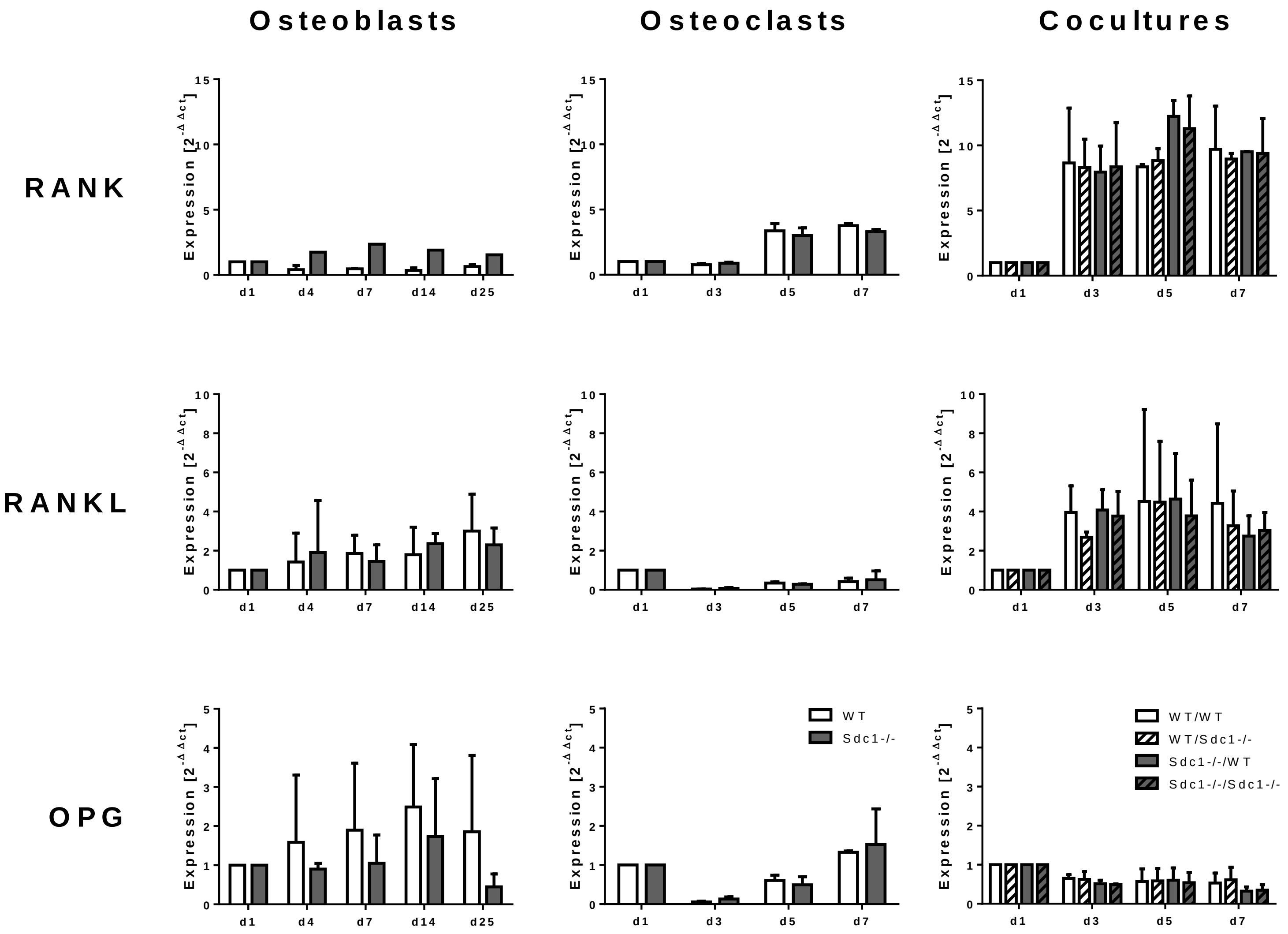

Supplemental figure 2

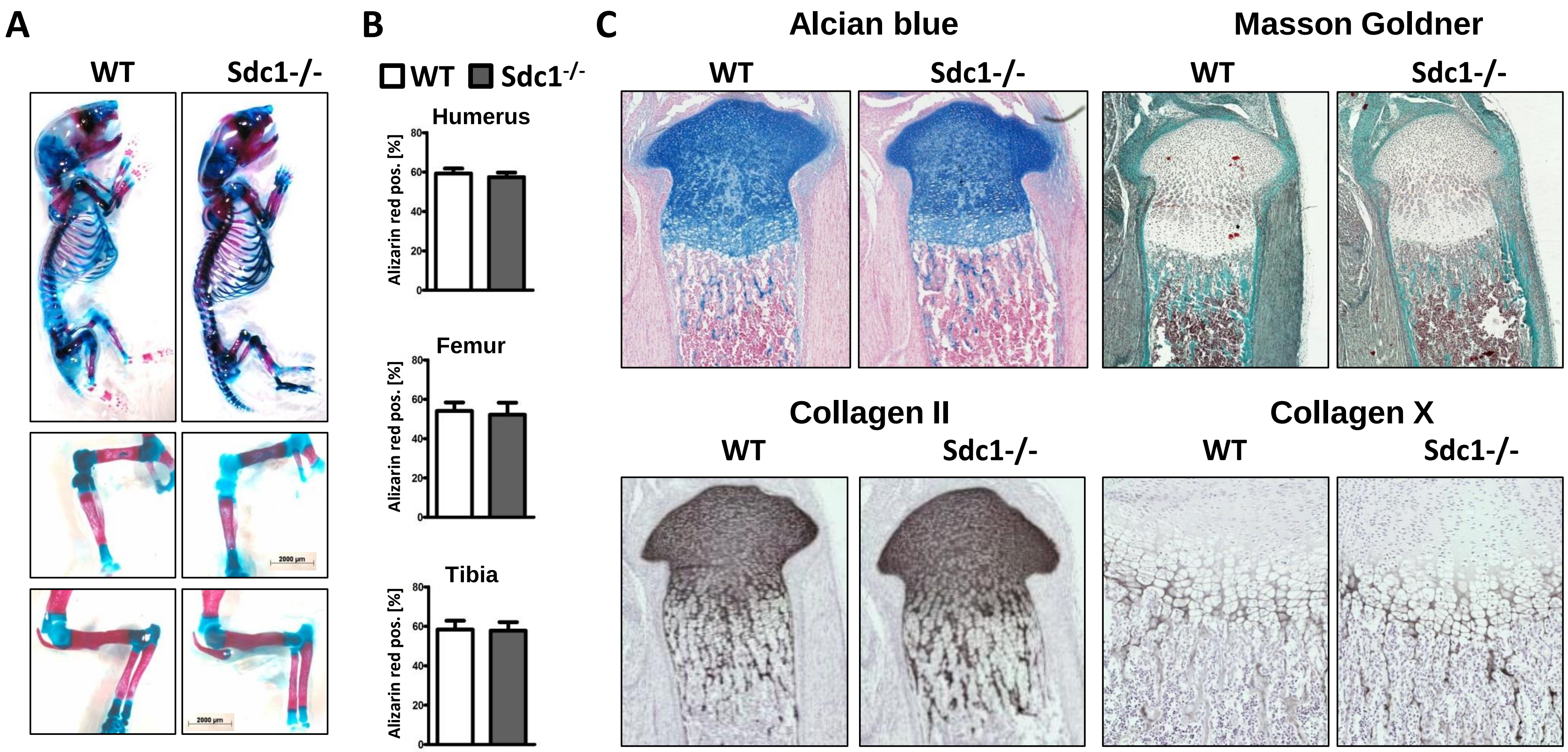

### Supplemental table 1

| Genotype | n | Age [month] | TV [mm³] | Whole BV/TV [%] | Trabecle BV/TV [%] | TbN [1/mm] | TbTh [µm] | TbS [µm] | CtTh [µm] |
|---|---|---|---|---|---|---|---|---|---|
| Wild type | 6 | 4 | 5.5 ±0.86 | 43.7±4.0 | 27.6±3.80 | 3.18±0.27 | 87.3±13.31 | 257±27.3 | 97.2±4.68 |
| Syndecan-1-/- | 9 | 4 | 3.5 ±0.79 | 43.6±3.37 | 28.91±3,6 | 3.31±0.25 | 87.3±6.41 | 252±15.2 | 95.3±3.84 |
| Wild type | 8 | 12 | 7.5± 0.49 | 39.5±2.3 | 26.7±3.27 | 2.93±0.30 | 89.4±4.49 | 307±28,1 | 95.8±1.61 |
| Syndecan-1-/- | 6 | 12 | 7.5± 0.74 | 37.6±3.71 | 24.4±3.40 | 2.76±0.38 | 86.8±2.18 | 301±21.3 | 94.1±2.3 |
| Wild type | 8 | 18 | 8.0± 0.93$^{\#}$ | 36.1±5.25 | 22.3±5.51 | 2.71±0.46 | 104.0±16.68 | 318±36.6 | 99.2±6.99 |
| Syndecan-1-/- | 10 | 18 | 7.9± 0.51$^{\#\#\#\#}$ | 33.8±2.89$^{\#\#\#}$ | 20.3±2.99$^{\#\#}$ | 2.30±0.16$^{\#\#\#\#}$ | 88.8±9.36 | 332±31.7$^{\#\#\#\#}$ | 92.1±2.6 |
